# S-MiRAGE: A quantitative, secreted RNA-based reporter of gene expression and cell persistence

**DOI:** 10.1101/325092

**Authors:** Kinshuk Mitra, William N. Feist, Simone Anfossi, Enrique Fuentes-Mattei, Maria Ines Almeida, Jean J. Kim, George A. Calin, Aryeh Warmflash

## Abstract

Non-destructive measurements of cell persistence and gene expression are crucial for longitudinal research studies and for prognostic assessment of cell therapies. Here we describe S-MiRAGE, a platform that utilizes small secreted RNA molecules as sensitive and quantitatively accurate reporters of cellular processes. We demonstrate the utility of S-MiRAGE by monitoring the differentiation status of human embryonic stem cells *in vitro*, and tumor growth in a mouse model *in vivo*.

---

A number of proteins are frequently used as genetically-encoded reporters including fluorescent proteins^(1)^, luciferase^(2)^, and Magnetic Resonance Imaging (MRI) or Positron Emission Tomography (PET)-detectable proteins^(3)^, with the choice of protein depending on the needs of the application. All of these systems suffer from limited multiplexability and poor signal to noise in challenging applications^(4)^, for example, when used as a reporter of weakly expressed endogenous genes. Motivated by recent successes in adopting circulating nucleic acids (cNAs) for clinical use^(5)^, we built a platform in which small expressed RNA molecules are used as reporters. A previous attempt to use secreted synthetic nucleic acids demonstrated the feasibility of this approach, but suffered from limited quantitative correlations with existing reporters and high background *in vivo*^(6)^. Overcoming these limitations could allow for the development of synthetic cNA-based reporters for a variety of research applications, and, in the future, for use as biomarkers^(6, 7)^ for medical interventions lacking quantitative metrics of success such as autologous therapies.

S-MiRAGE (Synthetic-<u>Mi</u>croRNA-like <u>R</u>eporters <u>A</u>ssaying <u>G</u>ene <u>E</u>xpression, hereafter referred to as MiRAGE) consists of a non-targeting microRNA (miR) placed downstream of a gene of interest (Fig. 1a). MicroRNAs encapsulated in extracellular vesicles (microvesicles and exosomes) or bound to protein (Ago2)^(8)^ and lipoprotein (HDL) are actively secreted by cells, and can be detected in the extracellular fluid^(9)^. The use of nucleic acids allows for multiplexing by simply changing the sequence, and sensitive detection via amplification by quantitative reverse transcription polymerase chain reaction (qRT-PCR). We placed the miR-30 backbone^(10)^ whose stem strands can be modified within a synthetic intron^(11)^ to reduce disruption of gene expression. We found that the MiRAGE reporter could be readily detected from the media of cultured cells, and that its expression level measured from media was the same whether placed at the N or C terminus of a protein-coding sequence (Supplementary Fig. 1a, b). We also found that probes against various miRs including a randomly generated sequence containing a previously described motif for exosomal sorting (ExoMotif)^(12)^ yielded some signal both in media from unmodified cells and mouse serum (Supplementary Fig. 1c). To reduce this background, we used qRT-PCR to test whether existing Taqman probes for zebrafish and *C. elegans* microRNAs produce background signals from mammalian biofluids (mouse serum and IP fluid). We selected two probes (cel-miR-2 and dre-miR-458, hereafter miR-2 and miR-458, respectively) that specifically detected their targets and exhibited no cross talk or background signal in mammalian biofluids for use in further studies.

**Figure 1.**
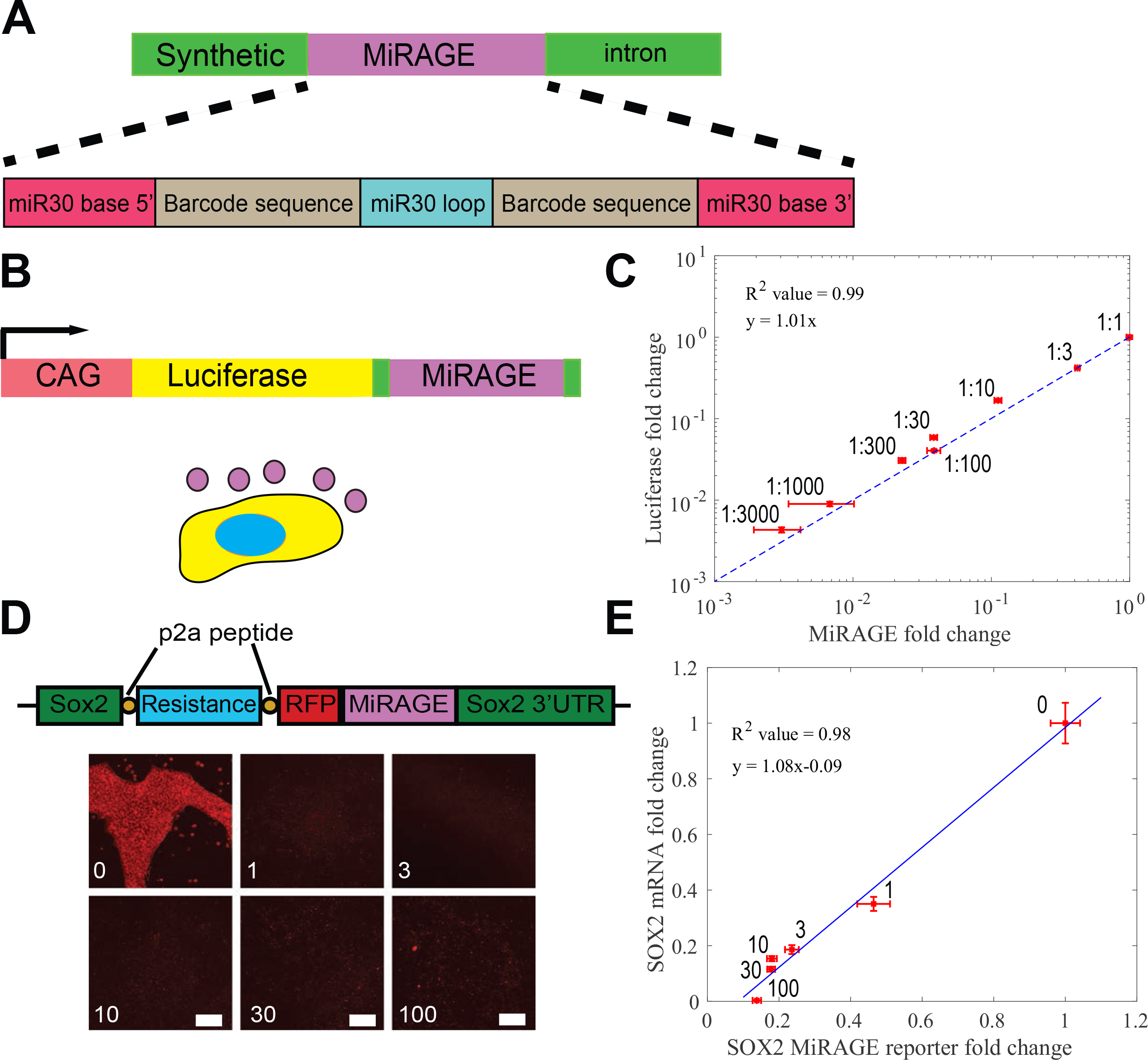
The MiRAGE system reports on cellular presence and endogenous gene expression. **a,** Schematic of the MiRAGE system based on a modified has-miR-30 sequence flanked by synthetic introns (green). The custom 22 or 23 nucleotide barcode sequence is labelled in brown. **b,** Schematic of a luciferase and MiRAGE expressing cell. MiRAGE is secreted by the cell while luciferase is retained intracellularly. **c,** Comparison of change in luciferase and MiRAGE reporter levels in media. 4T1-MiRAGE-luciferase cells were mixed with unmodified cells in varying proportions. Each data point is labeled with the ratio of reporter to wild-type cells; null hypothesis of slope of zero rejected at p<0.001. Fold changes are relative to 1:1 dilution. **d,** Schematic of modified SOX2 allele in the ESI017-SOX2-RFP-MiRAGE reporter cell line (top) and images of the RFP reporter upon treatment with the indicated concentrations of BMP4 for 4 days (bottom); scale bar 100 μm. **e,** Comparison of intracellular *SOX2* mRNA with the MiRAGE reporter assayed from culture media. Both the *SOX2* mRNA levels and the miR levels were determined by qPCR. mRNA is normalized to GAPDH while the MiRAGE reporter is normalized to miR-17. Data points are labeled by the dose of BMP4 (in ng/ml); null hypothesis of slope of zero rejected at p<0.001. Fold changes are as compared to a BMP4 dose of 0 ng/ml. In (c) and (e), each data represents averages and standard deviations across three biological replicates.

Luciferase is a commonly used reporter due to its sensitivity and wide dynamic range^(13)^. We placed a MiRAGE cassette within a synthetic intron at the 3’ end of firefly luciferase under the control of a constitutive promoter and used the ePiggyBac system to stably integrate this construct into mouse 4T1 cells (Fig. 1b). We mixed these cells with unlabeled 4T1 cells at varying ratios and observed a perfect linear relationship between the luciferase signal and MiRAGE levels detected by qRT-PCR on samples extracted from the culture media (Fig. 1c, R^2^=0.99). This linear relationship had a slope of one showing that the fold changes measured by MiRAGE and luciferase were identical over a large dynamic range (>10^3^).

We next evaluated whether our system can monitor endogenous gene expression. We integrated a MiRAGE reporter, RFP, and a blasticidin resistance gene at the C-terminus of the *SOX2* gene, a marker for pluripotency, in the human embryonic stem cell (hESC) line ESI017 using CRISPR-Cas9^(14)^ (Fig.1d). We determined that modifications to the wild-type *SOX2* allele occurred in less than 5% of cells heterozygous for the reporter (Supplementary Figure 2 a, b). We treated these reporter cells with varying concentrations of Bone Morphogenic Protein 4 (BMP4), causing differentiation to a trophectoderm-like fate, and loss of SOX2 expression^(15, 16)^. Fluorescent signal from the integrated RFP gene was completely lost at doses as low as 1 ng/ml (Fig. 1d), which mirrored the loss of SOX2 protein as measured by immunofluorescence (Supplementary Fig. 3). We compared the level of the MiRAGE reporter measured from either the media or the cells with the amount of *SOX2* mRNA in the same cells and found a nearly perfect correlation (Fig. 1e, R^2^=0.98 and Supplementary Fig. 4). The best correlation between cellular *SOX2* mRNA and MiRAGE was observed when the expression of the latter was normalized to that of a constitutively expressed microRNA, miR-17^(17)^, rather than an mRNA (GAPDH), potentially revealing variability in the total secretion of miRNA which can be accounted for by proper normalization (Supplementary Fig. 4). Thus, the MiRAGE reporter was a quantitative, non-destructive readout for the levels of SOX2 mRNA which varied more gradually with BMP4 levels than either the SOX2 protein or the RFP reporter.

We next used the MiRAGE-*SOX2* reporter to monitor reprogramming to pluripotency directly from the culture media. We differentiated ESI017-MiRAGE-S*OX2* cells into fibroblasts using an established protocol^(18)^, and confirmed differentiation via immunostaining that showed loss of the pluripotency marker SOX2 and upregulation of the fibroblast marker Vimentin (Supplementary Fig. 5). We then initiated reprogramming by over-expressing canonical reprogramming factors: OCT4, KLF4, SOX2 and c-MYC using a Sendai virus platform^(19)^. Imaging of the cells and analysis of MiRAGE levels in cell-culture media indicated that the expression of both reporter types (RFP and MiRAGE) fell below the threshold of detection during differentiation. Following the initiation of reprogramming, the reporters reappeared and gradually rose to the level in undifferentiated cells (Fig. 2a-b). We compared the MiRAGE reporter with *SOX2* mRNA at critical points in the differentiation and reprogramming process, and again found a nearly perfect correlation (Fig. 2c).

**Figure 2.**
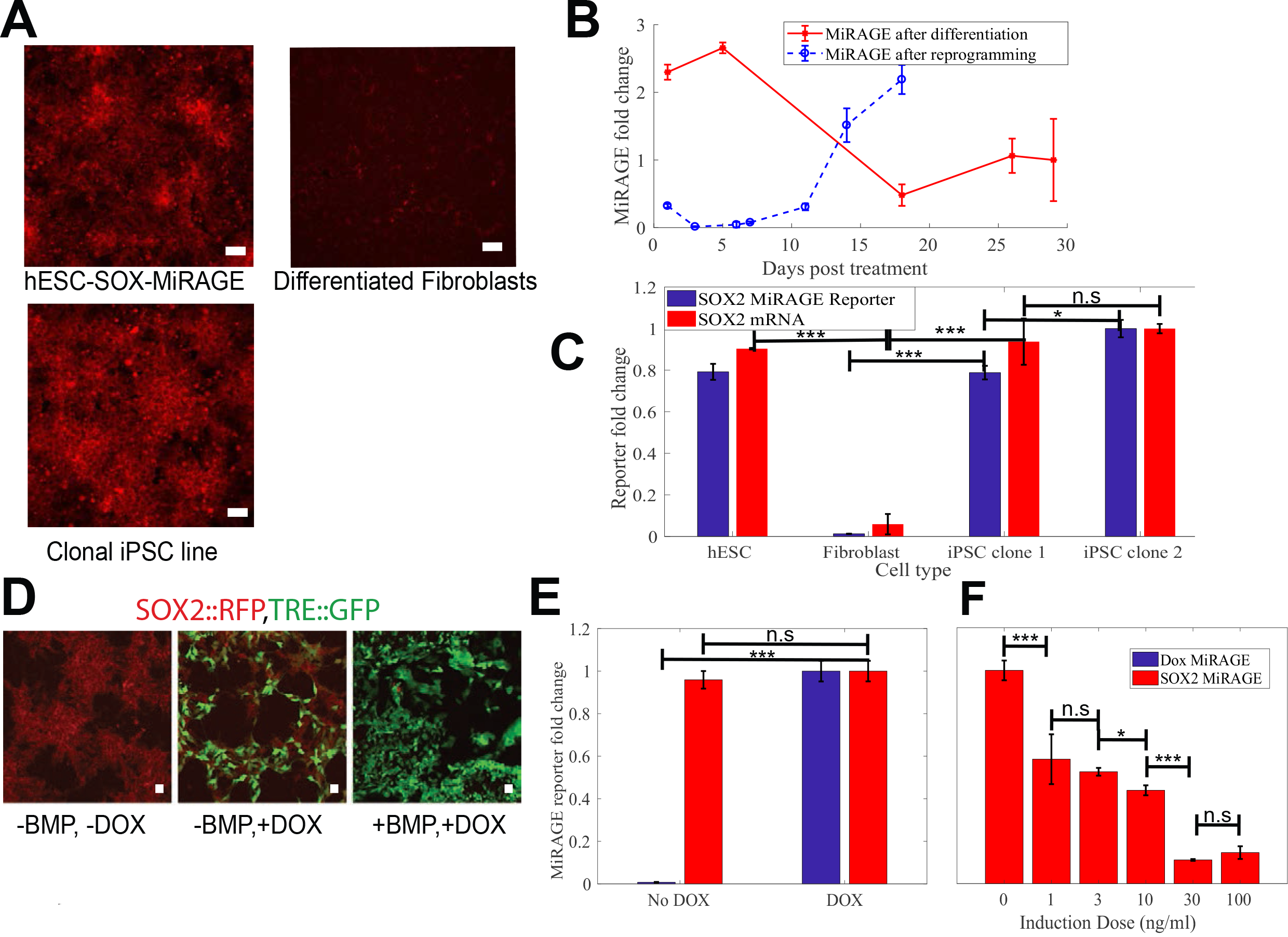
MiRAGE reports on reprogramming and allows for multiplexed monitoring of gene expression. **a,** Images of ESI017-SOX2-RFP-MiRAGE cells at the indicated points in the reprogramming and differentiation process; scale bar 50 μm. **b,** Expression level of the MiRAGE reporter in time during differentiation to fibroblasts and reprogramming assayed from the culture media by qPCR. MiRAGE levels were measured after beginning differentiation of hESCs into fibroblasts (day 0, differentiation) or after the removal of Sendai virus from cell culture during reprogramming (day 0, reprogramming). Fold changes are relative to the MiRAGE reading at the 28^th^ day post differentiation. **c,** Comparison of qPCR measurements of *SOX2* mRNA levels to MiRAGE levels in undifferentiated hESCs, differentiated fibroblasts, and 2 reprogrammed iPSC clones. Dataset is normalized to readings obtained from iPSC clone 2. **d,** Expression of fluorescent reporters of SOX2 and doxycycline with addition of either BMP4 or doxycycline; scale bar 20 μm. **e,** Multiplexed detection of MiRAGE reporters for doxycycline and *SOX2* mRNA from the same cells in the presence and absence of Dox. Each reporter is normalized to readings taken under Dox induction. **f,** SOX2 MiRAGE expression levels as a function of BMP4 dose in the presence of Dox. Fold changes are relative to the value at 0 BMP with Dox. Dox was used at a concentration of 100 ng/ml.

Current fluorescent and luminescent reporters also suffer from a limited capacity to multiplex due to spectral overlap. In contrast, we expected MiRAGE reporters with different stem sequences to be orthogonal. As proof of principle, ESI017-MiRAGE-SOX2 cells were engineered to express miR-2 and GFP in response to the addition of doxycycline (Fig. 2d). Adding doxycycline strongly induced the miR-2 MiRAGE reporter without affecting the expression of the miR-458 reporter of *SOX2* expression (Fig 2e). Similarly, the downregulation of the miR-458 upon addition of BMP was not affected by the doxycycline-induced upregulation of miR-2 (Fig. 2f).

A limited subset of reporter genes can be detected *in-vivo* due to the restricted ability to detect signal through deep tissue. We reasoned that MiRAGE, detected by a blood test, could function as an efficient reporter for applications not requiring spatial information and negate the need for expensive instrumentation. To test this hypothesis, we injected nude female mice intraperitoneally with HeYA8 ovarian cancer cells engineered to express luciferase and either MiRAGE miR-458 or MiRAGE miR-2 (Fig. 3a; 10 mice each group). Five of the miR-2 mice did not survive the experiment due to rapid cancer progression and were not included in the analysis, so that 15 mice were analyzed in total. At either second or fourth week after HeYA8 cell injection, blood and intraperitoneal (IP) fluid were harvested, the blood processed into plasma, and total small RNA extracted from both biofluids. We assayed all fluids for both miR-2 and miR-458 and found that the miR detected in both plasma and IP fluid (15/15 mice in both cases) matched that expressed in the cell line injected into the animal while the miR expressed by the non-injected cell line was never detected in the blood plasma (0/15 mice) and only in a small number of cases in the IP fluid (3/15 mice) (Fig. 3b, Supplementary Fig. 6a). Thus, MiRAGE is a sensitive and specific means to detect the presence of engineered cancer cells in vivo, particularly from the blood plasma.

**Figure 3.**
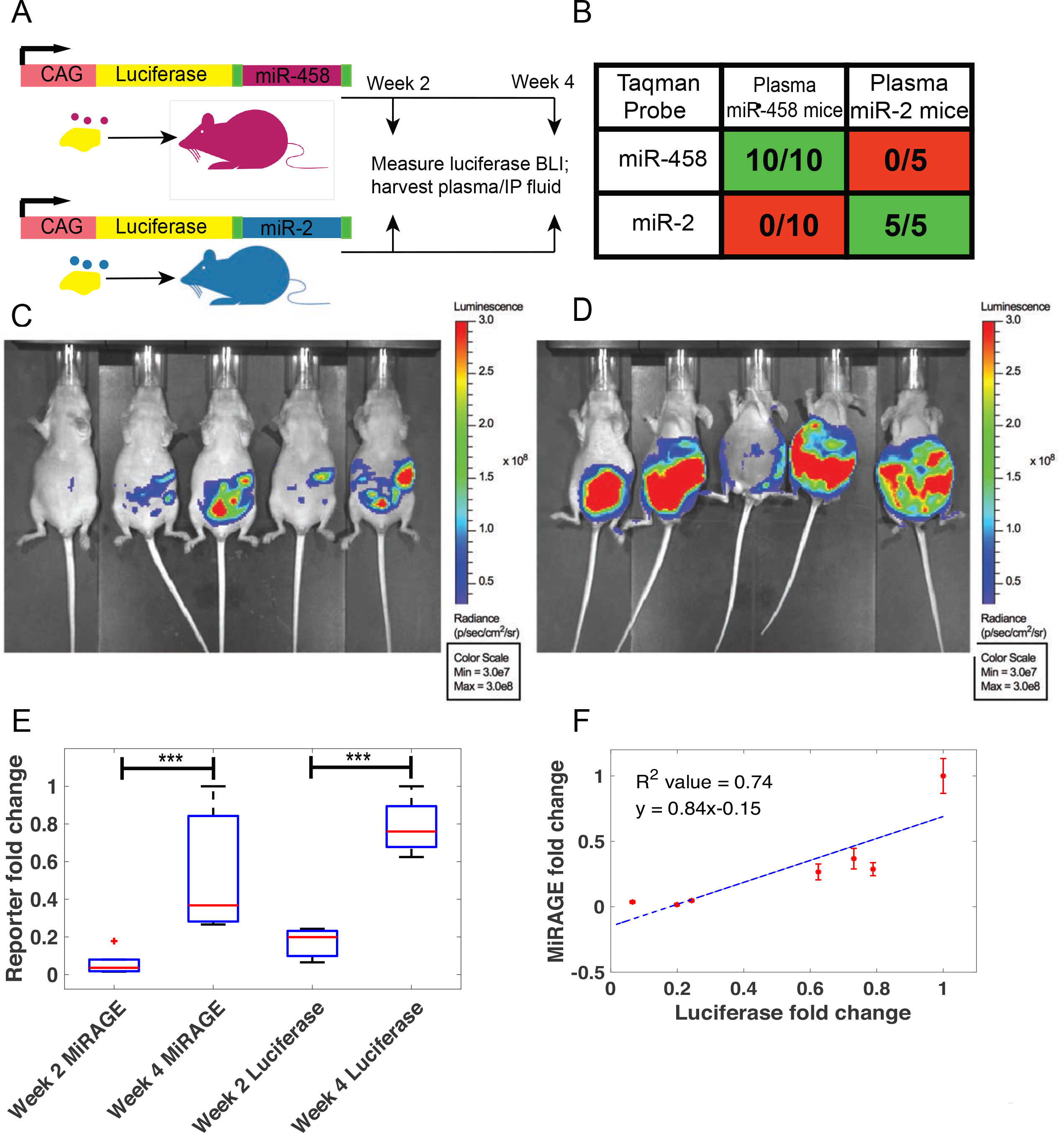
MiRAGE can be used to report on cell presence in vivo. **a,** Schematic of the experiment. **b** Table showing the fraction of mice in which the indicated probe yielded positive results from the indicated blood plasma sample. All successful detections represent samples with CT values of less than 35 for all three replicates, while all negative samples had CT values greater than 35 for all three replicates. **c, d,** Whole body bioluminescent images of miR-458 mice after week 2 (b) and week 4 (c) of tumor implantation. **e,** Boxplot of MiRAGE and luciferase signal across week 2 and week 4 miR-458 mice cohorts; 5 mice per boxplot group. Data for each reporter is normalized to the mouse with the highest expression of that reporter. **f,** Comparison of the level of circulating MiRAGE determined by qPCR from blood plasma with luciferase dosimeter signal detected for miR-458 mice. Each data point corresponds to one animal. We removed from consideration three mice whose IP fluid buildup interfered with the bioluminescent imaging; null hypothesis of slope of zero rejected at p<0.05. Datasets for MiRAGE and luciferase are normalized to their maximum values.

To determine whether the quantitative levels of the MiRAGE reporter can be used as readout of tumor growth, mice were also imaged for total whole-body bioluminescence after 2 and 4 weeks prior to biofluid collection (Fig. 3c, d). We found that the fold increase in MiRAGE signal from plasma was identical to that determined by luminescence (Fig. 3e). We compared MiRAGE signal from plasma to luciferase dosimeter values in individual mice from both week 2 and week 4 cohorts. We observed strong agreement between the two reporters with the MiRAGE signal increasing somewhat faster than the luminescence at high levels (Fig. 3f, R^2^=0.74). In contrast, whereas we did detect the MiRAGE reporter in the IP fluid, its levels did not correlate quantitatively with that of the luciferase, possible due to poor clearance from the IP compartment or dilution of MiRAGE by increased IP fluid accumulation (Supplementary Fig. 6b, c).

Our results here demonstrate proof of principle that a secreted miRNA reporter can be used as a quantitative, multiplexable readout of cell numbers or gene expression both in vitro and in vivo. Leveraging recent advances in methods for RNA detection^(20)^ will further improve the sensitivity of these measurements. Thus, MiRAGE will be a powerful tool for multiplexed non-destructive measurements and could potentially be used as a surrogate secreted biomarker when a natural one is absent.

## Acknowledgements

The authors thank Ali Brivanlou for the plasmid ePB-tta-puro-synthetic intron-shOCT4-RFP with the synthetic intron, and Alessandro Rosa for advice on using the synthetic intron. We also thank the flow cytometry core at MD Anderson, the Human Stem Cell Core at Baylor College of Medicine, and the members of the A.W lab for helpful discussions and feedback. This work was funded by grants to AW from the Cancer Prevention and Research Institute of Texas (CPRIT grant number RR140073) and the National Science Foundation (NSF grant MCB-1553228). Work in Dr. Calin’s laboratory is supported by National Institutes of Health (NIH/NCATS) grant UH3TR00943-01 through the NIH Common Fund, Office of Strategic Coordination (OSC), the NIH/NCI grant 1 R01 CA182905-01, a U54 grant – UPR/MDACC Partnership for Excellence in Cancer Research 2016 Pilot Project, a Team DOD (CA160445P1) grant, a Ladies Leukemia League grant, a CLL Moonshot Flagship project, a SINF 2017 grant, and the Estate of C. G. Johnson, Jr.

